# Evidence of functional long-range Wnt/Wg in the developing *Drosophila* wing epithelium

**DOI:** 10.1101/412627

**Authors:** Varun Chaudhary, Michael Boutros

**Author notes:** Addresses for correspondence: and.

## Abstract

Wnts are secreted proteins that regulate cell fate specification during development of all metazoans. Wnt proteins were proposed to spread over several cell diameters to activate signalling directly at a distance. In the *Drosophila* wing epithelium, an extracellular gradient of Wingless (Wg, the homolog of Wnt1) was observed extending over several cells away from producing cells. However, it was also recently shown that a membrane-tethered Neurotactin-Wg fusion protein (NRT-Wg) can rescue the loss-of endogenous Wg, leading to proper patterning of the wing. Therefore, whether Wg spreading is required for correct tissue patterning during development remains controversial and the functional range of wild-type Wg is unclear. Here, by capturing secreted Wg on distally located cells we show that the Wg gradient acts directly up to eleven cell distances. Cells located outside the reach of extracellular Wg depend on the Frizzled2 receptor to maintain target gene expression. We find that NRT-Wg is not restricted to the producing cells and propose that it can rescue signalling defects by perdurance in the receiving cells. These results provide insight into the mechanisms by which Wnt proteins mediate patterning of a rapidly growing tissue.

## INTRODUCTION

Wnts are lipid-modified secreted proteins conserved in all metazoans and play a pivotal role in development and many diseases (Nusse and Clevers, 2017; Zhan et al., 2017). The role of Wnt as a potential long-range signalling molecule has been controversially discussed as different methods to elucidate the range of extracellular Wnt and to manipulate gradient formation have generated contradictory paradigms for its range of action. On the one hand, direct and indirect visualization approaches can detect Wnt proteins up to several cell distances away from Wnt producing cells in *C. elegans* and *Drosophila* (Neumann and Cohen, 1997; Pani and Goldstein, 2018; Zecca et al., 1996). In addition, studies of Wnt protein carriers, for example exosomes and other vesicles, lipoprotein particles and proteinaceous co-factors (Beckett et al., 2013; Greco et al., 2001; Gross et al., 2012; Korkut et al., 2009; Luga et al., 2012; Mulligan et al., 2012; Neumann et al., 2009; Panáková et al., 2005) support a model whereby Wnt proteins can travel over long distances. Furthermore, a broad and graded expression of Wnt target genes with a localized Wnt expression domain has been observed during patterning of the mammalian primary body axis, displaying morphogen-like activities (Aulehla et al., 2003; Aulehla et al., 2008; Gao et al., 2011; Gavin et al., 1990; Kiecker and Niehrs, 2001; Niehrs, 2010). However, the long-range morphogen model for Wnt proteins has been challenged by recent studies, showing that juxtacrine Wnt signalling to the neighbouring cell by a membrane-tethered Wnt protein is sufficient for proper tissue patterning in *Drosophila* (Alexandre et al., 2014). Furthermore, a primarily short-range distribution of Wnt proteins is observed in the mouse intestinal crypts mediated by membrane association and cell division (Boutros and Niehrs, 2016; Farin et al., 2016), and Wnts are also found on the surface of cytonemes and filopodia (Huang and Kornberg, 2015; Mattes et al., 2018; Stanganello et al., 2015; Stanganello and Scholpp, 2016).

Arguably, the strongest competing models for the mode of action of Wnt proteins have been proposed based on experiments performed in the developing *Drosophila* wing epithelium – the wing imaginal disc. During early stages of wing discs development Wg is expressed broadly in the wing primordium, however, at later stages, it is restricted to two rows of cells at the dorso-ventral (DV) boundary (Couso et al., 1993; García-García et al., 1999; Ng et al., 1996; Williams et al., 1993). Previously, it was shown that Wg can spread over several cell distances to directly activate the expression of high-threshold targets, such as Senseless and low-threshold targets like Distalless (Dll) in a concentration dependent manner (Cadigan et al., 1998; Neumann and Cohen, 1997; Struhl and Basler, 1993; Zecca et al., 1996), although the exact functional range of Wg remained unknown. Clonal expression of Wg, but not the membrane-tethered NRT-Wg fusion protein, generated a gradient and activated target gene expression (Zecca et al., 1996). In contrast to this, a recent study showed that knock-in of NRT-Wg can almost completely replace endogenous Wg, supporting normal tissue patterning, albeit with growth defects (Alexandre et al., 2014), suggesting that Wg spreading is not necessary for normal patterning of wing imaginal discs. These competing models raise two fundamental questions; (1) what is the direct functional range of endogenous Wg protein, and (2) how is the membrane-tethered NRT-Wg able to generate similar wing patterning as endogenous Wg?

Understanding the functional range of Wg proteins for patterning the wing epithelium has so far been challenging due to the confounding effect from parallel tissue growth. Proliferation of cells leads to their movement away from the localized source of the extracellular protein gradient (Dekanty and Milan, 2011). While target gene expression in these cells would be expected to reduce gradually, it was, however, shown that expression of Wg target genes that require a low-level of signalling were in fact sustained in the absence of Wg (Piddini and Vincent, 2009). This suggests that mechanisms, although largely unknown, may exist to compensate for the decreasing concentration of Wg experienced by cells moving out of the Wg gradient.

In this study, we captured Wg proteins at a distance from the producing cells and measured the direct range of action of Wg gradient by generation of single cell clones expressing Frizzled2 (Fz2) receptor. By this means, we demonstrate that Wg can spread over at least 11 cells to activate signalling. Surprisingly, we also find that at the extracellular level membrane-tethered NRT-Wg is not restricted to the producing and adjacent receiving cells, but can also be detected further away. We propose that higher protein perdurance of NRT-Wg, perhaps due to the properties of the NRT protein, could contribute to its ability to pattern the wing imaginal disc. Furthermore, we show that cells that have moved out of the range of a functional Wg gradient then require the Fz2 receptor to maintain the expression of low-threshold Wg target genes. Altogether, our study demonstrates that Wg morphogen signalling in the growing epithelia is regulated by both long-range gradient-dependent and gradient-independent mechanisms.

## RESULTS

### Wg acts up to a distance of eleven cells

We first set out to investigate the direct range of a functional Wg gradient. Previous studies have shown that overexpression of Fz2 causes Wg stabilization at the cell surface (Bhanot et al., 1996; Cadigan et al., 1998; Piddini et al., 2005). We made use of this observation to probe how far from the DV boundary the Fz2 receptor could capture Wg protein and enhance Wg dependent signalling output. We generated single-cell Fz2 overexpression clones using the heat-shock inducible flip-out system. In order to control for the effect of cell proliferation on the accuracy of determining the Wg range, we induced the expression of Fz2 only at mid third instar larval stage and reduced the growth rate by keeping larvae at 18°C after the heat shock. By these means, we recovered single-cell clones of Fz2 (marked by nuclear GFP) in the late third instar wing disc (Figure 1A-E arrows and Figure S1). These GFP-marked cells showed accumulation of Wg (Figure 1C-C’) and a higher level of Dll expression (Figure 1D-D’) than surrounding control cells. We determined that these Wg-accumulating cells can be found up to 11 cell distances away from the DV boundary (Figure 1A’ and E’ yellow dots).

**Figure 1:**
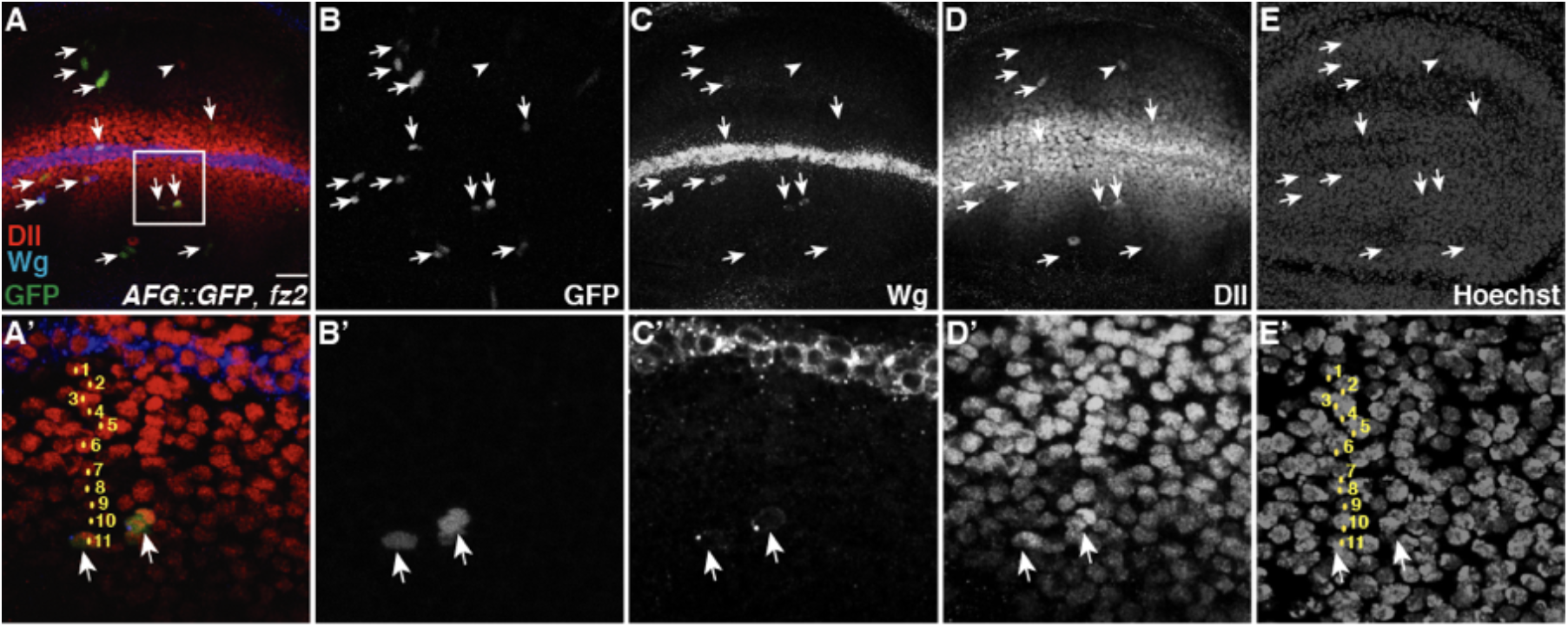
Wg can be detected up to 11 cell diameters from the Wg producing cells. **(A–E)** Single cell flip-out clones overexpressing Fz2 were generated in the discs (marked with GFP, arrows). 24 hours after the heat-shock to induce Fz2 expressing single-cell clones, total Wg staining shows clones with accumulation of Wg (C, images show projection of several confocal slices). Dll levels were modestly up-regulated in these GFP positive cells while very few cells also show Dll up-regulation without the apparent expression of GFP (B, arrowhead). **(A’-E’)** Area marked with white box in A was enlarged and a merge of 3 confocal slices was used to count the number of nuclei between the Fz2 overexpressing clone and Wg producing cells. GFP positive cells (A’ and B’ arrows) showing higher levels of Wg (C’) and Dll (D’) are at a distance of approximately 11 nuclei from the Wg producing cells (A’ and E’, nuclei marked with yellow dots). Scale bar 20 μm. N≥4 discs (see also Figure S1).

To further confirm these results, we generated single-cell clones overexpressing a GPI-linked N-terminal fragment of Fz2, which acts as a dominant negative. Consistent with the Fz2 overexpression, these single-cell clones accumulated Wg protein, but did not upregulate Dll expression (Figure S2A-D), suggesting that Dll activation required the formation of functional Wg-Fz2 complex. Taken together with previous observations that Wg expression at the late third instar stage is restricted to the DV boundary (Couso et al., 1993; García-García et al., 1999; Ng et al., 1996; Williams et al., 1993), these results suggest that endogenously secreted Wg can reach a distance of up to 11 cells to directly regulate transcriptional targets.

### Membrane-tethered Wg protein is detected in Wg receiving cells

The results presented above raises the question, how NRT-Wg, which is thought to be restricted to the expressing cells, can induce a similar patterning of the tissue as wild-type Wg. To address this question, we analyzed whether extracellular NRT-Wg protein remains restricted to the Wg expressing cells or if it can also be detected in cells that respond to Wg outside its normal expression domain. To this end, we performed extracellular Wg staining on third instar discs homozygous for *NRT-wg*. We then analysed the distribution of extracellular Wg by dividing the images into small regions of interests (ROI) (corresponding approximately to one row of cells) across the DV boundary and measured fluorescent intensity in each ROI (Figure 2A-A’). At this stage of disc development, it is expected that NRT-Wg, being membrane-tethered, will be restricted to the two stripe of *wg* expressing cells at DV boundary and the adjacent layer of receiving cell, however the extracellular NRT-Wg protein was detected over a broader domain covering several receiving cells (Figure 2A’). Furthermore, the levels of extracellular NRT-Wg protein were also much higher in the producing cells as compared to the endogenous Wg, which indicates that the NRT-Wg protein may be more stable than the endogenous Wg

**Figure 2:**
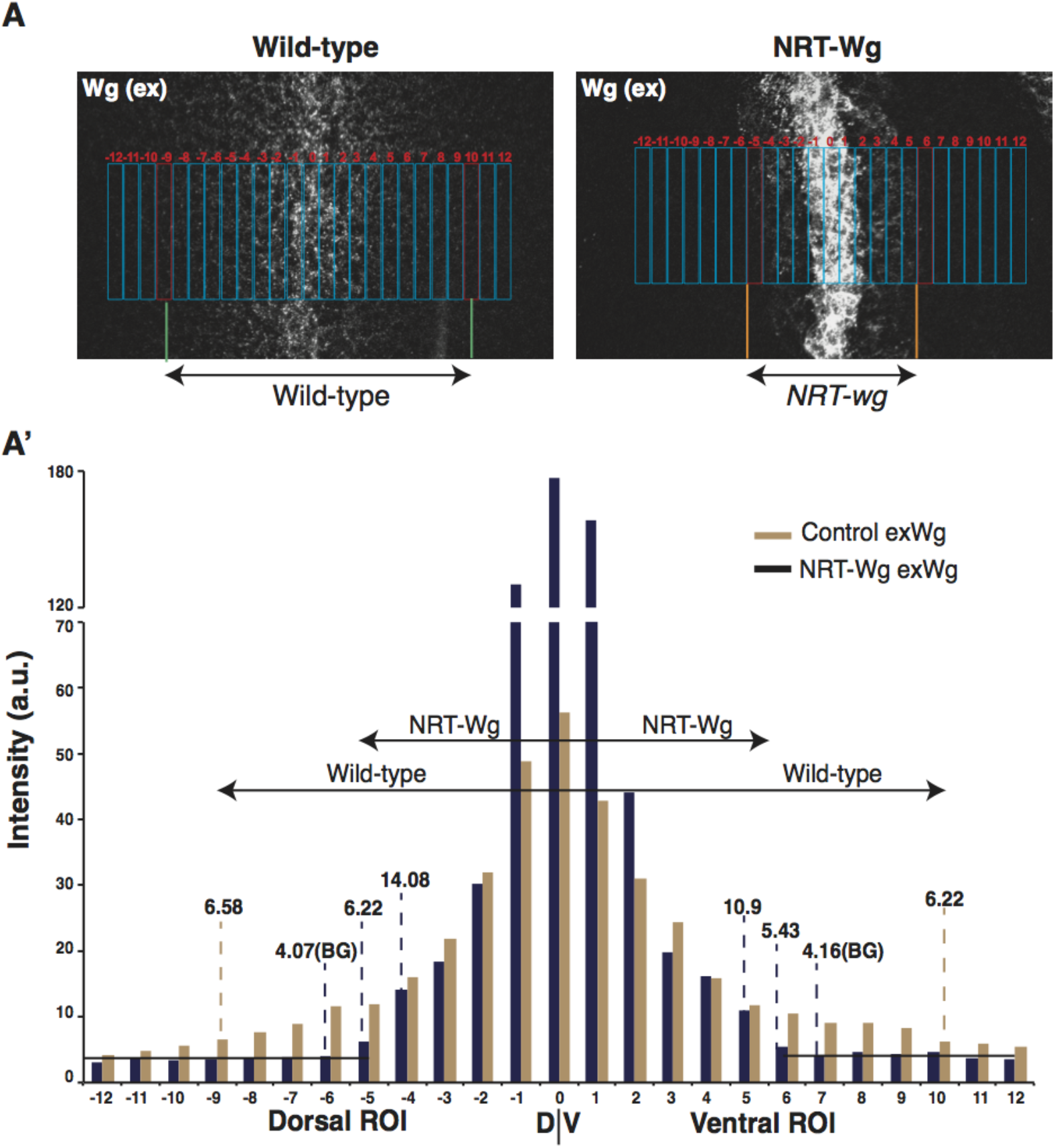
Comparison of extracellular Wg staining of wild-type and NRT-Wg wing imaginal discs. Quantification of the extracellular Wg in the homozygous *NRT-wg* or wild-type discs (quantification of discs shown in Fig. 3): (A) Distribution of extracellular Wg in the homozygous *NRT-wg* or wild-type (*w^1118^*) discs was divided into 25 rectangular ROI across the DV boundary (-12 to DV to 12), where each ROI represents approximately a single row of cells. (A’) Intensity was measured for each ROI separately and ROI on the either side of the DV boundary where the intensity was higher than the background signal (BG, approx. 4) were selected for determining the range of extracellular Wg on the *NRT-wg* discs (from segment −5 to between 5 and 6). ROI showing similar intensities were used for determining the range of extracellular Wg on wild-type discs (from region −9 to 10). Images were processed and analysed using ImageJ, under identical conditions.

**Figure 3:**
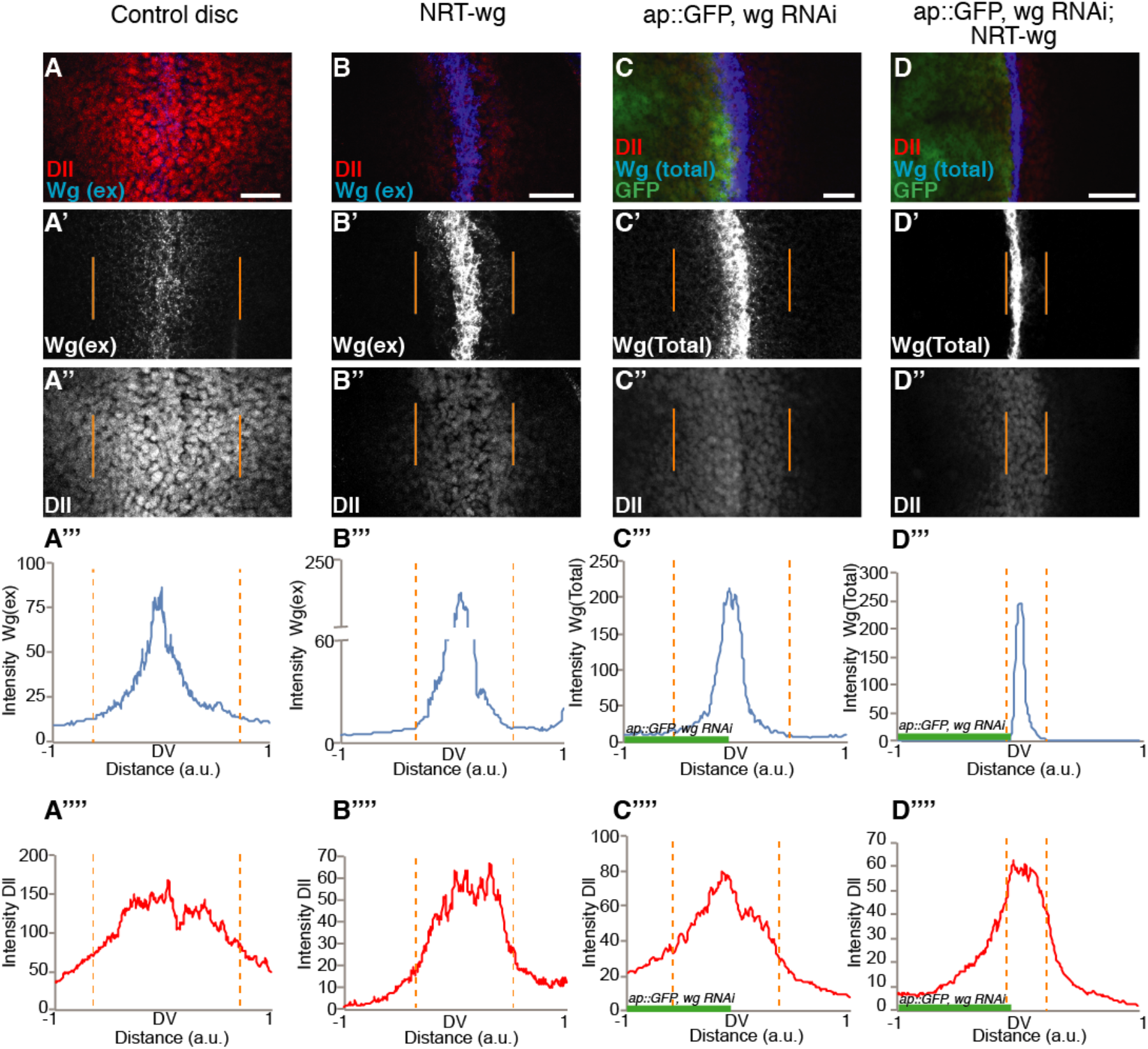
Presence of membrane-tethered Wg in the receiving cells rescues signalling. **(A-B)** Representative images of extracellular Wg and Dll staining on either wild-type **(A–A”)** or NRT-Wg homozygous **(B–B”)** discs imaged with the same confocal settings. Orange lines mark the extent of the region where extracellular Wg could be detected **(A”’, A”” and B”’, B””)** Graphs show intensity profile with x-axis of similar length as the shown images. **(C–D)** Total Wg and Dll staining on discs with depletion of Wg in dorsal compartment (left, marked by GFP) with *ap-Gal4, UAS-GFP/UAS-wg-RNAi* in either wild-type (C–C”) or NRT-Wg homozygous (D–D”) background (also see the diagrammatic representation below the panels, green cells represent *wg RNAi* expressing cells). **(C”’, C”” and D”’, D””)** Graphs show intensity profile with x-axis of similar length as the shown images.

Next, we analysed the range of Dll expression in these discs. We observed that Dll was expressed at higher levels in cells which were also positive for NRT-Wg, compared to cells where extracellular NRT-Wg was below detectable levels (Figure 3A”-B” and Figure S3A”-B”). This indicates that the expression of target genes in *NRT-wg* discs could be directly regulated by the presence of NRT-Wg protein on the surface of cells outside the normal Wg expression domain.

### Retention of membrane-tethered Wg in the receiving cells rescues signalling defects

We next asked whether the presence and effect of both endogenous ‘wild-type’ Wg and NRT-Wg in the receiving cells (no longer expressing Wg mRNA) requires their expression in these cells. To this end, we continually depleted Wg or NRT-Wg specifically in the dorsal compartment of the disc by expressing *wg* RNAi under the control of the *ap-Gal4* driver (Figure 3C-D and Figure S3C-D, left side of the disc also marked by GFP), leaving the ventral compartment and the ventral stripe of Wg producing cells unaffected. Considering first the depletion of Wg in a wild-type background where Wg is secreted from the ventral stripe of producing cells and spreads over several cells in both directions, we were able to detect Wg (total Wg staining) positive puncta in *wg* RNAi expressing cells of the dorsal compartment (Figure 3C, C’ and C”’ and Figure S3C-C’, left side of the disc also marked by GFP). Accordingly, Dll expression in the dorsal compartment was similar to the ventral control compartment (Figure 3C”, C”” and Figure S3C”). In contrast, expression of *wg* RNAi in the dorsal compartment of *NRT-wg* homozygous discs led to a strong reduction of total NRT-Wg levels (Figure 3D, D’ and D”’ and Figure S3D-D’). Consequently, this led to a strong reduction in Dll expression in the dorsal compartment (Figure 3D”,D”” and Figure S3D”). These experiments suggest that unlike ‘wild-type’ Wg, the presence of NRT-Wg in the receiving cells is a result of its earlier expression and higher protein perdurance in these cells. Therefore, longer retention of NRT-Wg protein in the receiving cells is required for the expression of Wg target genes.

### Fz2 receptor activity maintains expression of long-range Wg target genes

Since Dll is expressed beyond 11 cells from the Wg expression domain, we next asked how Dll expression can be maintained in the absence of extracellular Wg. One way of maintaining Wnt signalling could be a modulation of the expression of feedback regulators, which can provide either positive or negative input at various levels of the pathway. To this end, we turned our attention to the Fz2 receptor, which is a positive regulator of the pathway and transcriptionally repressed by Wg signalling (Cadigan et al., 1998). Reducing Wg function in discs during late larval stages causes up-regulation of Fz2 mRNA (Cadigan et al., 1998), whereas Dll expression remained unaffected (Piddini and Vincent, 2009). Therefore, we hypothesize that Dll levels could be maintained by the increased amount of Fz2 receptors in the cell.

First, we confirmed the up-regulation of Fz2 at the protein level using antibody staining. Blocking Wg secretion by depletion of the Wnt cargo-receptor Evenness interrupted (Evi; also known as Wntless) (Bartscherer et al., 2006; Bänziger et al., 2006; Goodman et al., 2006) in the posterior compartment led to a modest increase in Fz2 protein levels (Figure S4A-A’, compare arrowhead to arrow). Interestingly, in these cells Dll remained expressed (Figure S4A”). Next, we asked whether the increase in Fz2 protein levels upon blocking Wg secretion supports maintenance of Dll expression. To address this, we generated *fz2* loss-of-function clones in discs expressing *evi* RNAi under the control of either *wg-Gal4* or *en-Gal4*, leading to a loss of secreted Wg either in the entire disc or the posterior compartment, respectively. As shown in Figure 4A-F (for *wg-Gal4>evi RNAi*) and Figure S4B-C (for *en-Gal4>evi RNAi*), we observed a significant loss of Dll expression (Figure 4D, arrows and Figure S4B”, C” arrows) in *fz2* mutant clones (marked by loss of GFP) compared to the neighbouring GFP positive control tissue. Therefore, these data demonstrate that in the absence of the extracellular Wg gradient upon Evi depletion, cells up-regulate Fz2 levels, which is required for the maintenance of Dll expression.

**Figure 4:**
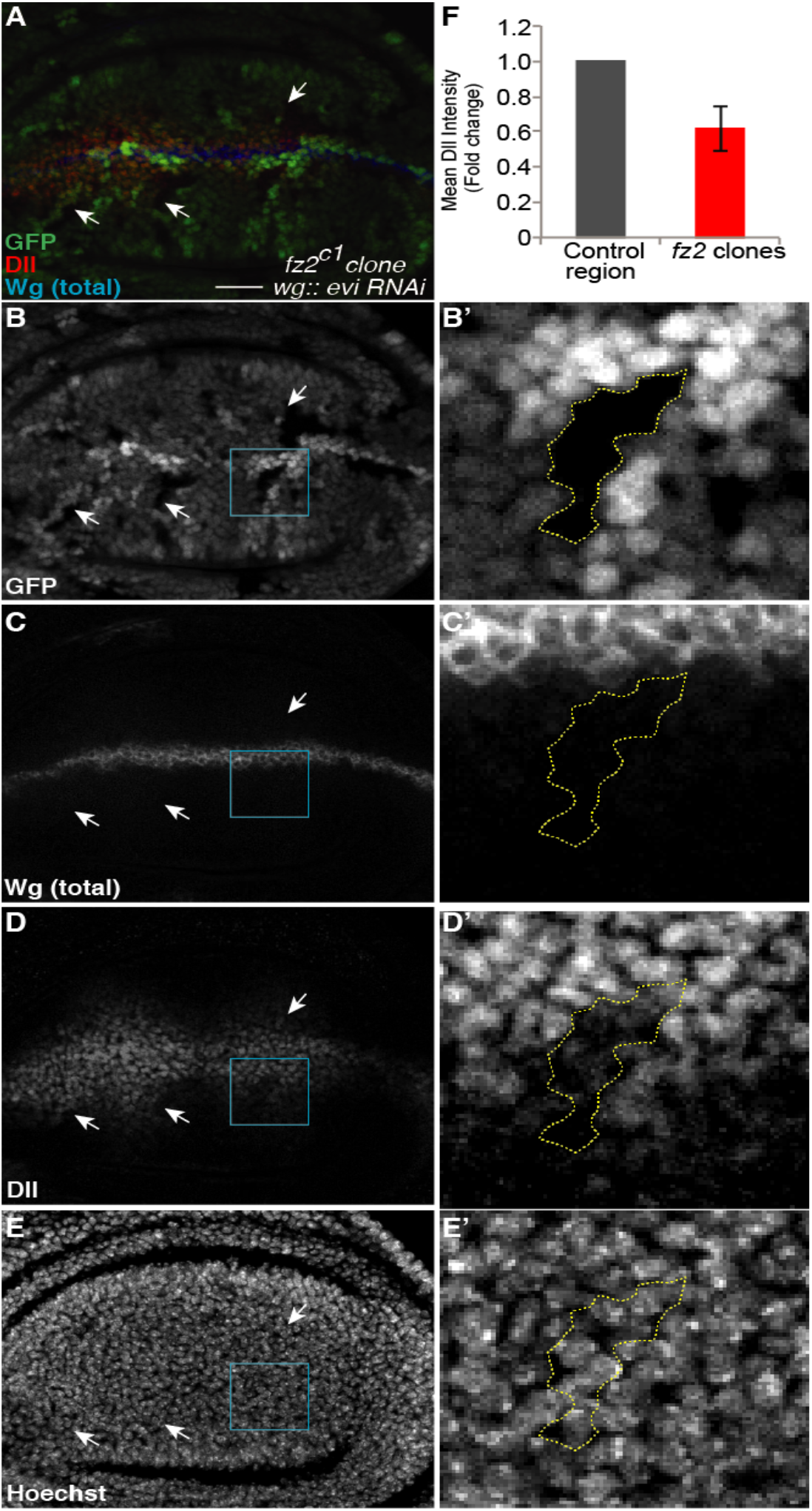
Fz2 is required for the maintenance of Dll expression in the absence of Wg secretion. Genotype of disc shown in this figure is *Ubx-FLP/+; wg-Gal4/UAS-evi-RNAi; fz2^c1^, π, FRT2A/ubi-GFP FRT2A*. **(A–E)** *fz2^c1^* clones (marked by absence of GFP) were generated in discs also expressing *evi-RNAi* leading to accumulation of Wg in the producing cells due to block in Wg se–cretion (C). Dll expression is reduced inside the *fz2^c1^* clones (D, arrows). **(B’-E’)** show enlarged region outlined in B–E. Dll expression is reduced inside the clone close to the receiving cells (**C**’ and **D**’). Hoechst staining shows presence of nuclei inside the clone as in neighbouring control cells (E’). **(F)** Quantification of Dll intensity with in the *fz2^c1^* clones and the neighbouring control region at the same distance from the DV boundary (N=12 clones). Scale bar 20 μm.

Fz2 acts redundantly with Frizzled1 (Fz1) in canonical Wnt signalling (Bhanot et al., 1999; Bhat, 1998; Chen and Struhl, 1999; Kennerdell and Carthew, 1998; Muller et al., 1999). We next asked whether Fz2, which is expressed at higher levels in cells away from the DV boundary has a non-redundant role with Fz1 in maintaining Dll expression. To test this, we analysed Dll expression in *fz2* mutant clones either close or away from DV boundary in otherwise wild-type discs (Figure S4D-F). Under this condition, *fz2* clones, which were close to the DV boundary (marked by white box in Figure S4D) showed no effect on Dll levels (Figure S4E-E”’), as previously shown (Chen and Struhl, 1999). However, in contrast, *fz2* clones further away from the DV boundary (marked by yellow box in Figure S4D) showed reduced levels of Dll expression compared to control cells at the same distance (Figure S4F-F”’, compare arrows marking clones to arrowhead marking control cell). However, the *fz1* clones showed no difference in Dll levels at both short and long-range (Figure S5A-D). Notably, in wing discs where Wg secretion was blocked, remaining Dll expression was reduced in *fz2* mutant clones irrespective of the location of the clone from the DV boundary (Figure 4B’-E’), supporting the conclusion that Wg-independent expression of Dll is Fz2 dependent.

### Higher levels of Fz2 protein in NRT-Wg discs may contribute to its rescue effects

Based on these results we hypothesized that in NRT-Wg discs, low level of Dll expression observed in cells outside the reach of NRT-Wg would be due to maintenance by Fz2 receptor. In addition, we find higher levels of Fz2 receptors in NRT-Wg homozygous discs compared to the control (Figure 5A and B). Collectively, these data demonstrate that while short-range signalling is Wg dependent and can be mediated by either Fz2 or Fz1, the long-range expression of Dll is Wg independent and regulated only by Fz2. Moreover, this indicates that Fz2-mediated maintenance signalling may play an important role in the overall survival of NRT-Wg animals.

**Figure 5:**
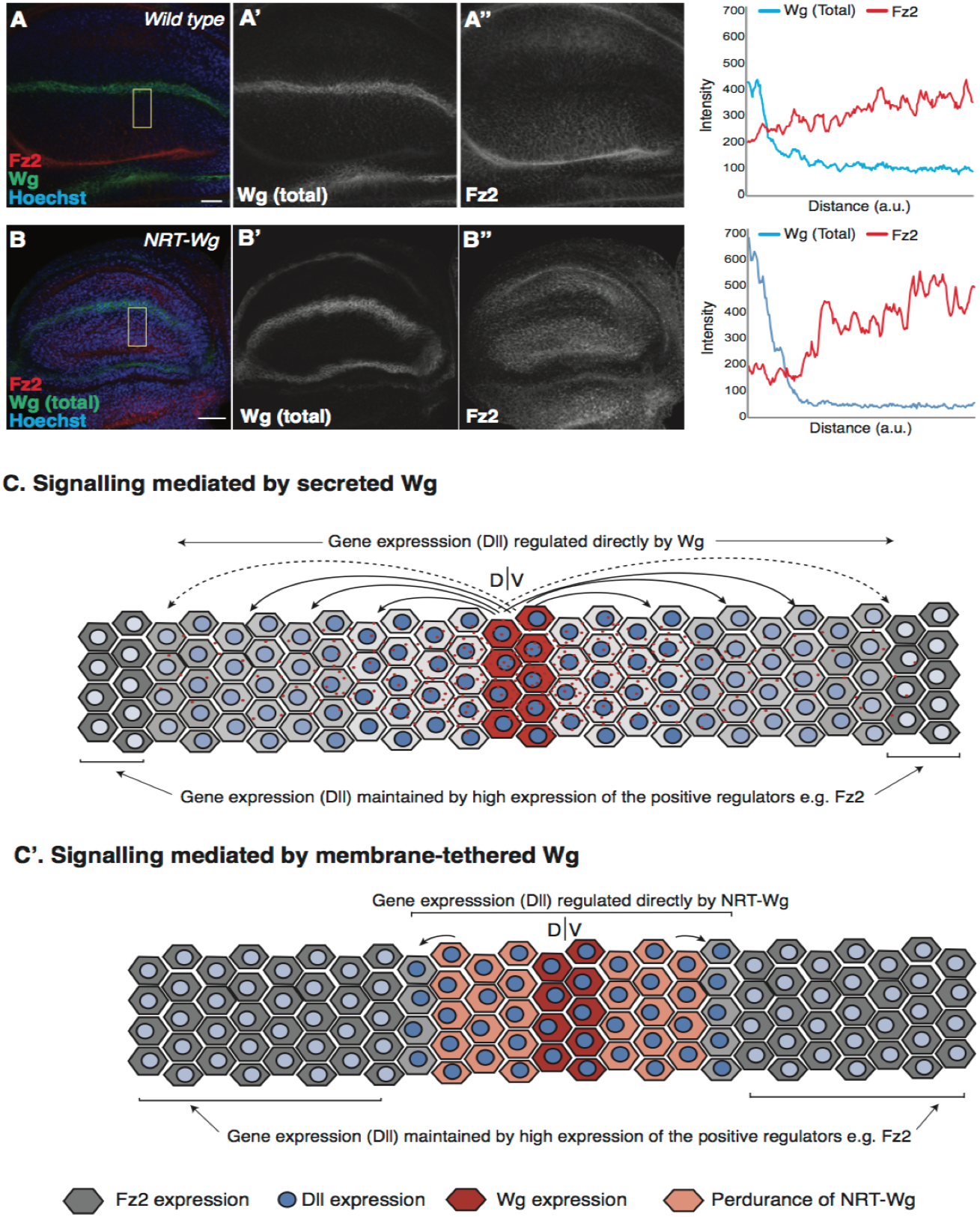
Increased levels of Fz2 in receiving cells may support survival of NRT-Wg flies. (A–B) Representative images of Fz2 and Wg staining on either wild-type (A–A”) or NRT-Wg homozygous (B–B”) discs imaged with the same confocal settings. Graphs on the right-side show intensity profile across the yellow box in A and B. (C – C’) Model representing the mechanism of signalling via secreted Wg which can act up to a distance of at least 11 cells. Cell beyond the range of secreted Wg can maintain Dll expression by upregulating positive regulators for example, Fz2 receptor. Whereas the expression of membrane-tethered NRT-Wg mediates signalling up to a few cell distances. Scale bar 20 μm.

## DISCUSSION

How secreted proteins influence patterning decisions at a distance has been a longstanding and fundamental question in developmental biology (Gierer and Meinhardt, 1972). It has been proposed that signalling molecules can act as morphogens by being secreted from a localized source and form gradients across a tissue to induce patterning decisions in a concentration dependant manner (Gierer and Meinhardt, 1972; Rogers and Schier, 2011; Wolpert, 1969, 1971). However, the biochemical nature of these gradients and their functional ranges are in many cases not well understood. It also remains largely unresolved how cells moving away from the source of such ‘morphogen’ signalling factors maintain their identity in a rapidly growing tissue.

In this study, we provide experimental evidence that Wg/Wnt signalling in target cells of the developing wing epithelium is regulated by the combined inputs from (1) a long-range Wg activity detected at a 11 cell distance and (2) the maintenance of Wg/Wnt signalling by the Fz2 receptor in cells exposed to gradually decreasing levels of Wg.

### Determining the signalling range of Wg in the developing *Drosophila* wing

Although almost four decades have passed since the discovery of Wnt, the mechanism by which Wnt proteins mediate long-range effects still remains poorly understood. Contradictory models proposing either a direct long-range spreading or juxtacrine signalling of Wnt proteins to the immediate neighbouring cell have been discussed (Pani and Goldstein, 2018; Neumann and Cohen, 1997; Zecca et al., 1996; Alexandre et al., 2014; Bartscherer et al., 2008; Morata and Struhl, 2014). In this study, by overexpression of the Fz2 receptor in single cell clones, we demonstrate that Wg can be captured at a distance of at least 11 cells from its origin. Single cell cell clones were generated at a late stage of third instar development when Wg expression is restricted to a narrow stripe at the DV boundary. These experiments also showed that capturing of Wg by Fz2 leads to a cell-autonomous increase in Dll expression, indicating that captured extracellular Wg in functional. Taken together, our results suggest that secreted Wg is active up to a distance of 11 cells in developing wing imaginal discs, concordant with its visualization in the extracellular space.

### Higher perdurance of membrane-tethered Wg can compensates for its inability to act at a distance

If Wg proteins are able to travel long-distance and mediate signalling, then how can a wild-type, secreted Wg protein be functionally replaced by a membrane-tethered version? Our experiments provide several lines of evidence that point towards a higher perdurance of NRT-Wg protein as compared wild-type Wg, as also previously discussed by Morata and Struhl (2014). The higher perdurance of NRT-Wg leads to the presentation of the ligand in cells that under wild-type conditions would not express, but receive a Wg signal (Figure 5C). First, immunofluorescence staining of extracellular NRT-Wg protein showed that it is not limited to Wg producing cells but can also be detected in ‘receiving’ cells which, presumably as a consequence, express high-levels of the Wg target gene Dll. Second, when we depleted NRT-Wg by RNAi in the dorsal compartment of *NRT-wg* homozygous discs, this led to a complete removal of NRT-Wg dorsally, while its expression appeared normal in the ventral compartment. In contrast, in ‘wild-type’ Wg discs, we observed a gradient of Wg and a similarly reduced expression pattern in both the RNAi knockdown and control compartments. While it is possible that NRT-Wg protein is transferred from one cell to other via cell division, as recently suggested for mammalian Wnt3 (Farin et al., 2016), our results suggest that the wild-type Wg protein gradient observed in wing discs is generated by secretion and spreading and not due to cell division and cell lineage-based expression. Interestingly, a recent study showed that NRT-Wg is unable to replace the endogenous Wg for proper patterning of the malpighian tubules (Beaven and Denholm, 2018), supporting model whereby a rescue effect of NRT-Wg observed in tissues such as the wing imaginal may depends on the tissue specific compensatory mechanisms, for example maintenance of signalling.

### Maintenance of long-range low-threshold target genes by Fz2

While our results show that target gene expression at a distance of 11 cells can be under the direct control of a Wg gradient, how signalling activity of Wg beyond this range is established remained largely unresolved. Previous studies have suggested that the expression of Dll can be maintained after initiation by auto-regulatory mechanisms via cis-regulatory elements in a Wg independent manner (Estella et al., 2008; Galindo et al., 2011; Piddini and Vincent, 2009). However, the enhancers identified were found to be active mainly in the leg imaginal discs and did not explain the expression of Dll observed at long-range in the wing discs. In this study, we show that the Fz2 receptor, which is transcriptionally repressed by Wg signalling, is up-regulated upon either block of Wg secretion or in NRT-Wg discs in cells where Dll expression is maintained. Removal of Fz2 from these cells led to further reduction of Dll levels, indicating that Fz2 is required for the maintenance of Dll expression in the absence of continuous input from extracellular Wg.

Furthermore, *fz2* mutant clones generated in otherwise wild-type discs showed reduced levels of Dll in clones found only at long-range and not at short-range. Fz1 unlike Fz2 is uniformly expressed in the discs, therefore loss of Fz2 at short-range would be rescued by Wg dependent signalling via Fz1. However, Fz1 is unable to rescue long-range Dll expression in Fz2 clones indicating that this expression of Dll is not regulated directly by extracellular Wg. This suggests that cells at long-range have moved out of the functional Wg gradient, which led to an increase in Fz2 expression that further maintains Dll expression (Figure 5C and C’). Therefore, our data reveals a new role for the Fz2 receptor in maintaining Wnt signalling. While the importance of this maintenance of signalling for tissue development remains to studied, we have observed that growth and patterning in NRT-Wg flies is further worsened by removal of Fz2 (data not shown), which suggest that maintenance may play a role in normal tissue development.

Feedback regulation in gradient signalling is a common mechanism used in a variety of growing tissues by various other signalling molecules for example in the case of Dpp (Ben-Zvi et al., 2011; Vuilleumier et al., 2010). Positive regulators are transcriptionally repressed while the negative-feedback genes are transcriptionally activated. This allows cells moving out of the functional range of the Wg gradient to gradually reduce the levels of negative regulators, for example a decoy receptor Frizzled3 (Sato et al., 1999) and increase the amount of positive regulators, for example Fz2 (Cadigan et al., 1998) and the Wg co-receptor Arrow (Wehrli et al., 2000). Like a balancing pole used for tight-rope walking, this underlying mechanism in a rapidly growing tissue enables cells to balance signalling at different positions along the Wg concentration gradient.

### Conclusions

In summary, our data supports a model whereby the endogenous Wg protein can travel over distances of several cells to induce a Wnt signalling cascade. While the underlying biochemical mechanisms – such as secretory vesicles, proteinaceous particles or filopodia – remain unknown, our data indicate that developing tissues retain a high degree of plasticity to respond to changes in extracellular cues with multiple, interacting mechanisms. Maintenance and other compensatory mechanisms may play an important role to buffer fluctuations resulting in the proper patterning of the wing epithelium. It will be interesting to understand the mechanism of receptor-mediated maintenance of Wg signalling in more details and to identify additional maintenance mechanisms. This study also indicates that genome-engineered variants of proteins that influence its stability might lead to unexpected compensatory mechanisms *in vivo*.

## MATERIALS AND METHODS

### Drosophila genetics

The following *Drosophila* stocks were used: *wg-GAL4* (2^nd^ chr., a gift from S. Cohen), *ap-GAL4* 2^nd^ chr., (Montagne et al., 1999), *en-GAL4, UAS-GFP* 2^nd^ chr. (Thompson and Cohen, 2006). *hs-Flp* on 1^st^ chr. and *UAS-GFP* in 3^rd^ chr. were gifted by A. Teleman. *fz2^C1^, ri, FRT2A* (Chen and Struhl, 1999), *fz^p21^* is a functional null allele (Jones et al., 1996) gifted by D. Strutt, *ubi-GFP FRT2A* (A. Gould), *NRT-HA-Wg (Baena-Lopez et al., 2013), evi^2^* (Bartscherer et al., 2006). *UAS-Fz2* (BDSC #41794) UAS-Fz2(ECD)-GPI (dominant negative Fz2) (BDSC #44221), ubi-GFP FRT80 (BDSC #1620) and *Actin5C-FRT-CD2-FRT-Gal4* (BDSC #4779) lines were obtained from Bloomington Stock Center (Bloomington, Indiana, USA). *UAS-wg-RNAi* (GD-ID 13352) and *UAS-evi-RNAi* (KK-ID 103812) lines were obtained from Vienna *Drosophila* RNAi Center. All crosses were reared on standard culture medium at 25°C except where specifically mentioned.

### Antibodies

The following antibodies were used: Mouse Anti-Wg (4D4s, obtained from Developmental System Hybridoma Bank) used 1:5 for extracellular and 1:50 for total staining, Rabbit-anti-Dll 1:200 (a gift from S. Carroll). Fluorescent secondary antibodies used were Alexa-405, Alexa-488, Alexa-594 and Alexa-647 (Invitrogen) at 1:300 dilutions.

### Immunostainings, microscopy and image analysis

Extracellular Wg staining was performed as described previously (Strigini and Cohen,2000). Wing discs were mounted in Vectashield. Staining and microscopy conditions for discs used for comparisons were identical. Images were taken with a Leica SP5 confocal and were analyzed using ImageJ software (Schneider et al., 2012). Plot analysis and intensity measurements were performed on raw data processed with ImageJ. Separate channel images were assembled using Adobe Photoshop CS5.1.

### Heat shock induction of ‘flip-out’ clones

*hs-FLP/Actin-FRT-CD2-Stop-FRT-Gal4; UAS-Fz2/UAS-GFP* flip-out’ clones were generated by heat shocking mid third-instar larvae for 30 minutes at 37°C. Larvae were then shifted to 18°C and were dissected 24 hours later.

### Drosophila genotypes

The following genotypes were used in this study:

Figure 1, Figure S1: *y w hs-FLP/Actin-FRT-Stop-FRT-Gal4; UAS-Fz2/UAS-GFP*

Figure S2A-D: *y w hs-FLP/Actin-FRT-CD2-Stop-FRT-Gal4; UAS-Fz2-ECD-GPI/ UAS-GFP*

Figure 2A (left panel), Figure 3A, Figure S3A, Figure 5A: *w1118* (wild-type control)

Figure 2A (right panel), Figure 3B, Figure S3B, Figure 5B: *NRT-wg/NRT-wg*

Figure 3C, Figure S3C: *ap-Gal4/+; UAS-GFP/UAS-wg-RNAi*

Figure 3D, Figure S3D: *ap-Gal4, NRT-wg/NRT-wg; UAS-GFP/UAS-wg-RNA*

Figure 4A-E: *y w Ubx-FLP/+; wg-Gal4/UAS-evi-RNAi(KK); fz2^C1^, ri, FRT2A/ubi-GFPFRT2A*

Figure S4A: *en-Gal4, UAS-GFP/UAS-evi-RNAi(KK)*

Figure S4B-C: *y w Ubx-FLP/+; en-Gal4/UAS-evi-RNAi(KK); fz2^C1^, ri, FRT2A/ubi-GFP FRT2A*

Figure S4D-F: *y w Ubx-FLP/+; +/+; fz2^C1^, ri, FRT2A/ubi-GFP FRT2A*

Figure S5: *y w Ubx-FLP/+; +/+; fz^p21^FRT80/ubi-GFP FRT80*

## Supporting information

## ACKNOWLEDGEMENTS

We are grateful to Fillip Port, Maja Funk and Bojana Pavlocic for comments on the manuscript and members of the Boutros laboratory and David Strutt for helpful discussions. We would like to thank G. Struhl, S. Carroll, J.P. Vincent, A. Teleman, the Bloomington stock center and VDRC for *Drosophila* strains and reagents. We are also grateful for support from the DKFZ Light Microscopy Core Facility and the IISER Bhopal Microscopy Facility. Work in the laboratory of M.B. was supported by a DFG grant on Wnt signalling.

## REFERENCES

Alexandre, C., Baena-Lopez, A., and Vincent, J.P. (2014). Patterning and growth control by membrane-tethered Wingless. Nature 505, 180–185.

Aulehla, A., Wehrle, C., Brand-Saberi, B., Kemler, R., Gossler, A., Kanzler, B., and Herrmann, B.G. (2003). Wnt3a plays a major role in the segmentation clock controlling somitogenesis. Dev Cell 4, 395–406.

Aulehla, A., Wiegraebe, W., Baubet, V., Wahl, M.B., Deng, C., Taketo, M., Lewandoski, M., and Pourquié, O. (2008). A beta-catenin gradient links the clock and wavefront systems in mouse embryo segmentation. Nat Cell Biol 10, 186–193.

Baena-Lopez, L.A., Alexandre, C., Mitchell, A., Pasakarnis, L., and Vincent, J.P. (2013). Accelerated homologous recombination and subsequent genome modification in Drosophila. Development 140, 4818–4825.

Bartscherer, K., Pelte, N., Ingelfinger, D., and Boutros, M. (2006). Secretion of Wnt ligands requires Evi, a conserved transmembrane protein. Cell 125, 523–533.

Beaven, R., and Denholm, B. (2018). Release and spread of Wingless is required to pattern the proximo-distal axis of Drosophila renal tubules. Elife 7

Beckett, K., Monier, S., Palmer, L., Alexandre, C., Green, H., Bonneil, E., Raposo, G., Thibault, P., Le Borgne, R., and Vincent, J.P. (2013). Drosophila S2 cells secrete wingless on exosome-like vesicles but the wingless gradient forms independently of exosomes. Traffic 14, 82–96.

Ben-Zvi, D., Pyrowolakis, G., Barkai, N., and Shilo, B.Z. (2011). Expansion-repression mechanism for scaling the Dpp activation gradient in Drosophila wing imaginal discs. Curr Biol 21, 1391–1396.

Bhanot, P., Brink, M., Samos, C.H., Hsieh, J.C., Wang, Y., Macke, J.P., Andrew, D., Nathans, J., and Nusse, R. (1996). A new member of the frizzled family from Drosophila functions as a Wingless receptor. Nature 382, 225–230.

Bhanot, P., Fish, M., Jemison, J.A., Nusse, R., Nathans, J., and Cadigan, K.M. (1999). Frizzled and Dfrizzled-2 function as redundant receptors for Wingless during Drosophila embryonic development. Development 126, 4175–4186.

Bhat, K.M. (1998). frizzled and frizzled 2 play a partially redundant role in wingless signaling and have similar requirements to wingless in neurogenesis. Cell 95, 1027–1036.

Boutros, M., and Niehrs, C. (2016). Sticking Around: Short-Range Activity of Wnt Ligands. Dev Cell 36, 485–486.

Bänziger, C., Soldini, D., Schütt, C., Zipperlen, P., Hausmann, G., and Basler, K. (2006). Wntless, a conserved membrane protein dedicated to the secretion of Wnt proteins from signaling cells. Cell 125, 509–522.

Cadigan, K.M., Fish, M.P., Rulifson, E.J., and Nusse, R. (1998). Wingless repression of Drosophila frizzled 2 expression shapes the Wingless morphogen gradient in the wing. Cell 93, 767–777.

Chen, C.M., and Struhl, G. (1999). Wingless transduction by the Frizzled and Frizzled2 proteins of Drosophila. Development 126, 5441–5452.

Couso, J.P., Bate, M., and Martínez-Arias, A. (1993). A wingless-dependent polar coordinate system in Drosophila imaginal discs. Science 259, 484–489.

Dekanty, A., and Milán, M. (2011). The interplay between morphogens and tissue growth. EMBO Rep 12, 1003–1010.

Estella, C., McKay, D.J., and Mann, R.S. (2008). Molecular integration of wingless, decapentaplegic, and autoregulatory inputs into Distalless during Drosophila leg development. Dev Cell 14, 86–96.

Farin, H.F., Jordens, I., Mosa, M.H., Basak, O., Korving, J., Tauriello, D.V., de Punder, K., Angers, S., Peters, P.J., Maurice, M.M., et al. (2016). Visualization of a short-range Wnt gradient in the intestinal stem-cell niche. Nature 530, 340–343.

Galindo, M.I., Fernández-Garza, D., Phillips, R., and Couso, J.P. (2011). Control of Distal-less expression in the Drosophila appendages by functional 3’ enhancers. Dev Biol 353, 396–410.

Gao, B., Song, H., Bishop, K., Elliot, G., Garrett, L., English, M.A., Andre, P., Robinson, J., Sood, R., Minami, Y., et al. (2011). Wnt signaling gradients establish planar cell polarity by inducing Vangl2 phosphorylation through Ror2. Dev Cell 20, 163–176.

García-García, M.J., Ramain, P., Simpson, P., and Modolell, J. (1999). Different contributions of pannier and wingless to the patterning of the dorsal mesothorax of Drosophila. Development 126, 3523–3532.

Gavin, B.J., McMahon, J.A., and McMahon, A.P. (1990). Expression of multiple novel Wnt-1/int-1-related genes during fetal and adult mouse development. Genes Dev 4, 2319–2332.

Gierer, A., and Meinhardt, H. (1972). A theory of biological pattern formation. Kybernetik 12, 30–39.

Goodman, R.M., Thombre, S., Firtina, Z., Gray, D., Betts, D., Roebuck, J., Spana, E.P., and Selva, E.M. (2006). Sprinter: a novel transmembrane protein required for Wg secretion and signaling. Development 133, 4901–4911.

Greco, V., Hannus, M., and Eaton, S. (2001). Argosomes: a potential vehicle for the spread of morphogens through epithelia. Cell 106, 633–645.

Gross, J.C., Chaudhary, V., Bartscherer, K., and Boutros, M. (2012). Active Wnt proteins are secreted on exosomes. Nat Cell Biol 14, 1036–1045.

Huang, H., and Kornberg, T.B. (2015). Myoblast cytonemes mediate Wg signaling from the wing imaginal disc and Delta-Notch signaling to the air sac primordium. Elife 4, e06114.

Jones, K.H., Liu, J., and Adler, P.N. (1996). Molecular analysis of EMS-induced frizzled mutations in Drosophila melanogaster. Genetics 142, 205–215.

Kennerdell, J.R., and Carthew, R.W. (1998). Use of dsRNA-mediated genetic interference to demonstrate that frizzled and frizzled 2 act in the wingless pathway. Cell 95, 1017–1026.

Kiecker, C., and Niehrs, C. (2001). A morphogen gradient of Wnt/beta-catenin signalling regulates anteroposterior neural patterning in Xenopus. Development 128, 4189–4201.

Korkut, C., Ataman, B., Ramachandran, P., Ashley, J., Barria, R., Gherbesi, N., and Budnik, V. (2009). Trans-synaptic transmission of vesicular Wnt signals through Evi/Wntless. Cell 139, 393–404.

Luga, V., Zhang, L., Viloria-Petit, A.M., Ogunjimi, A.A., Inanlou, M.R., Chiu, E., Buchanan, M., Hosein, A.N., Basik, M., and Wrana, J.L. (2012). Exosomes mediate stromal mobilization of autocrine Wnt-PCP signaling in breast cancer cell migration. Cell 151, 1542–1556.

Mattes, B., Dang, Y., Greicius, G., Kaufmann, L.T., Prunsche, B., Rosenbauer, J., Stegmaier, J., Mikut, R., Özbek, S., Nienhaus, G.U., et al. (2018). Wnt/PCP controls spreading of Wnt/β-catenin signals by cytonemes in vertebrates. Elife 7.

Montagne, J., Stewart, M.J., Stocker, H., Hafen, E., Kozma, S.C., and Thomas, G. (1999). Drosophila S6 kinase: a regulator of cell size. Science 285, 2126–2129.

Morata, G., and Struhl, G. (2014). Developmental biology: Tethered wings. Nature 505, 162–163.

Mulligan, K.A., Fuerer, C., Ching, W., Fish, M., Willert, K., and Nusse, R. (2012). Secreted Wingless-interacting molecule (Swim) promotes long-range signaling by maintaining Wingless solubility. Proc Natl Acad Sci U S A 109, 370–377.

Müller, H.A., Samanta, R., and Wieschaus, E. (1999). Wingless signaling in the Drosophila embryo: zygotic requirements and the role of the frizzled genes. Development 126, 577–586.

Neumann, C.J., and Cohen, S.M. (1997). Long-range action of Wingless organizes the dorsal-ventral axis of the Drosophila wing. Development 124, 871–880.

Neumann, S., Coudreuse, D.Y., van der Westhuyzen, D.R., Eckhardt, E.R., Korswagen, H.C., Schmitz, G., and Sprong, H. (2009). Mammalian Wnt3a is released on lipoprotein particles. Traffic 10, 334–343.

Ng, M., Diaz-Benjumea, F.J., Vincent, J.P., Wu, J., and Cohen, S.M. (1996). Specification of the wing by localized expression of wingless protein. Nature 381, 316–318.

Niehrs, C. (2010). On growth and form: a Cartesian coordinate system of Wnt and BMP signaling specifies bilaterian body axes. Development 137, 845–857.

Nusse, R., and Clevers, H. (2017). Wnt/β-Catenin Signaling, Disease, and Emerging Therapeutic Modalities. Cell 169, 985–999.

Pani, A.M., and Goldstein, B. (2018). Direct visualization of a native Wnt. Elife 7.

Panáková, D., Sprong, H., Marois, E., Thiele, C., and Eaton, S. (2005). Lipoprotein particles are required for Hedgehog and Wingless signalling. Nature 435, 58–65.

Pfeiffer, S., Alexandre, C., Calleja, M., and Vincent, J.P. (2000). The progeny of wingless-expressing cells deliver the signal at a distance in Drosophila embryos. Curr Biol 10, 321–324.

Piddini, E., Marshall, F., Dubois, L., Hirst, E., and Vincent, J.P. (2005). Arrow (LRP6) and Frizzled2 cooperate to degrade Wingless in Drosophila imaginal discs. Development 132, 5479–5489.

Piddini, E., and Vincent, J.P. (2009). Interpretation of the wingless gradient requires signaling-induced self-inhibition. Cell 136, 296–307.

Rogers, K.W., and Schier, A.F. (2011). Morphogen gradients: from generation to interpretation. Annu Rev Cell Dev Biol 27, 377–407.

Sato, A., Kojima, T., Ui-Tei, K., Miyata, Y., and Saigo, K. (1999). Dfrizzled-3, a new Drosophila Wnt receptor, acting as an attenuator of Wingless signaling in wingless hypomorphic mutants. Development 126, 4421–4430.

Schneider, C.A., Rasband, W.S., and Eliceiri, K.W. (2012). NIH Image to ImageJ: 25 years of image analysis. Nat Methods 9, 671–675.

Serralbo, O., and Marcelle, C. (2014). Migrating cells mediate long-range WNT signaling. Development 141, 2057–2063.

Stanganello, E., Hagemann, A.I., Mattes, B., Sinner, C., Meyen, D., Weber, S., Schug, A., Raz, E., and Scholpp, S. (2015). Filopodia-based Wnt transport during vertebrate tissue patterning. Nat Commun 6, 5846.

Stanganello, E., and Scholpp, S. (2016). Role of cytonemes in Wnt transport. J Cell Sci 129, 665–672.

Strigini, M., and Cohen, S.M. (2000). Wingless gradient formation in the Drosophila wing. Curr Biol 10, 293–300.

Struhl, G., and Basler, K. (1993). Organizing activity of wingless protein in Drosophila. Cell 72, 527–540.

Thompson, B.J., and Cohen, S.M. (2006). The Hippo pathway regulates the bantam microRNA to control cell proliferation and apoptosis in Drosophila. Cell 126, 767–774.

Vuilleumier, R., Springhorn, A., Patterson, L., Koidl, S., Hammerschmidt, M., Affolter, M., and Pyrowolakis, G. (2010). Control of Dpp morphogen signalling by a secreted feedback regulator. Nat Cell Biol 12, 611–617.

Wehrli, M., Dougan, S.T., Caldwell, K., O’Keefe, L., Schwartz, S., Vaizel-Ohayon, D., Schejter, E., Tomlinson, A., and DiNardo, S. (2000). arrow encodes an LDL-receptor-related protein essential for Wingless signalling. Nature 407, 527–530.

Williams, J.A., Paddock, S.W., and Carroll, S.B. (1993). Pattern formation in a secondary field: a hierarchy of regulatory genes subdivides the developing Drosophila wing disc into discrete subregions. Development 117, 571–584.

Wolpert, L. (1969). Positional information and the spatial pattern of cellular differentiation. J Theor Biol 25, 1–47.

Wolpert, L. (1971). Positional information and pattern formation. Curr Top Dev Biol 6, 183–224.

Zecca, M., Basler, K., and Struhl, G. (1996). Direct and long-range action of a wingless morphogen gradient. Cell 87, 833–844.

Zhan, T., Rindtorff, N., and Boutros, M. (2017). Wnt signaling in cancer. Oncogene 36, 1461–1473.

