## Supplementary material for "Evidence of functional long-range Wnt/Wg in the developing *Drosophila* wing epithelium"

### SUPPLEMENTAL FIGURES

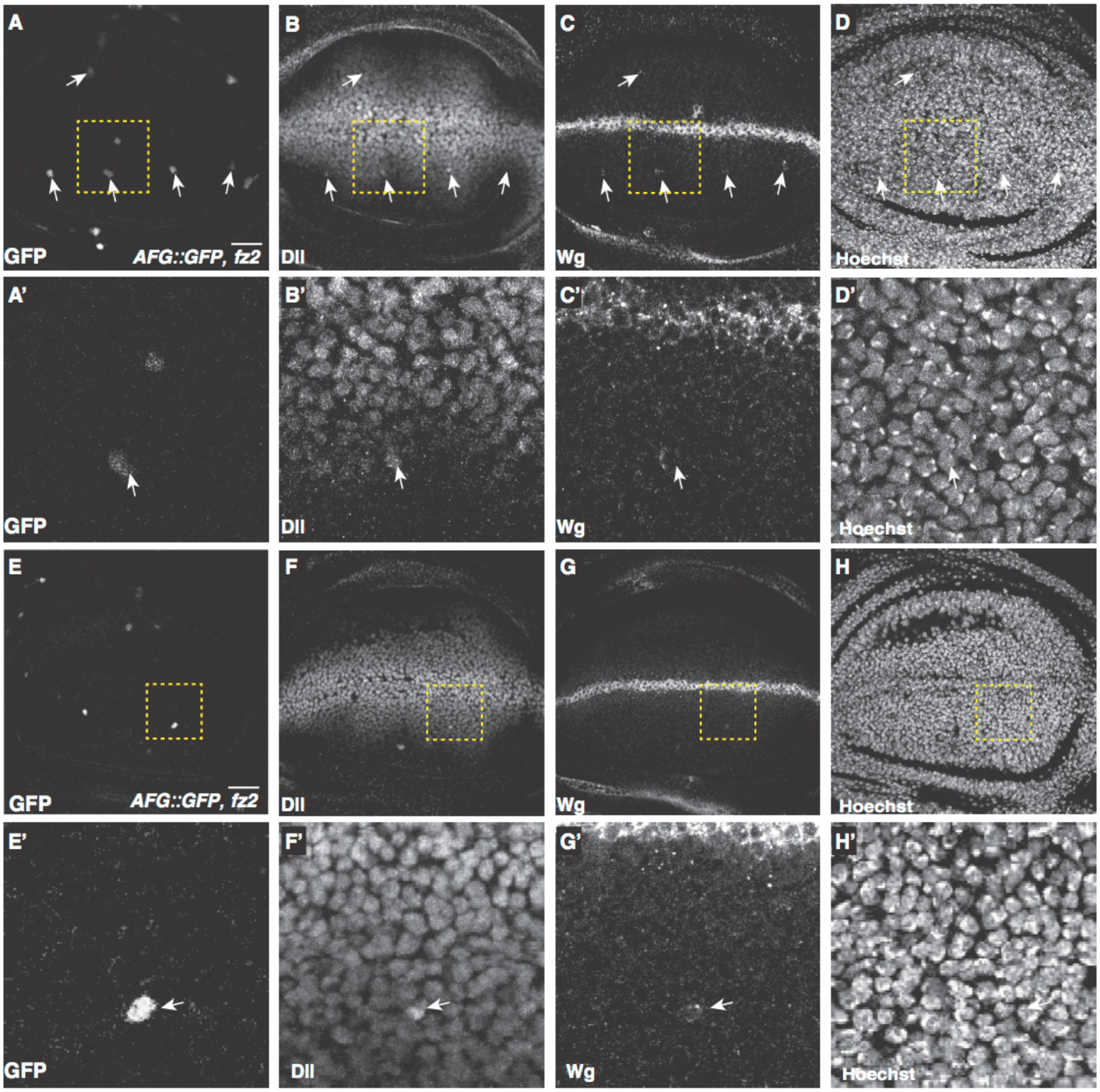


**Figure S1: Single-cell clones expressing Fz2 show Wg accumulation and enhanced signalling.** To supplement Figure 1, panels provide further two examples (A-D and E-H) of Wg accumulation and Dll upregulation in single-cell clones overexpressing Fz2. Scale bar 20 µm.

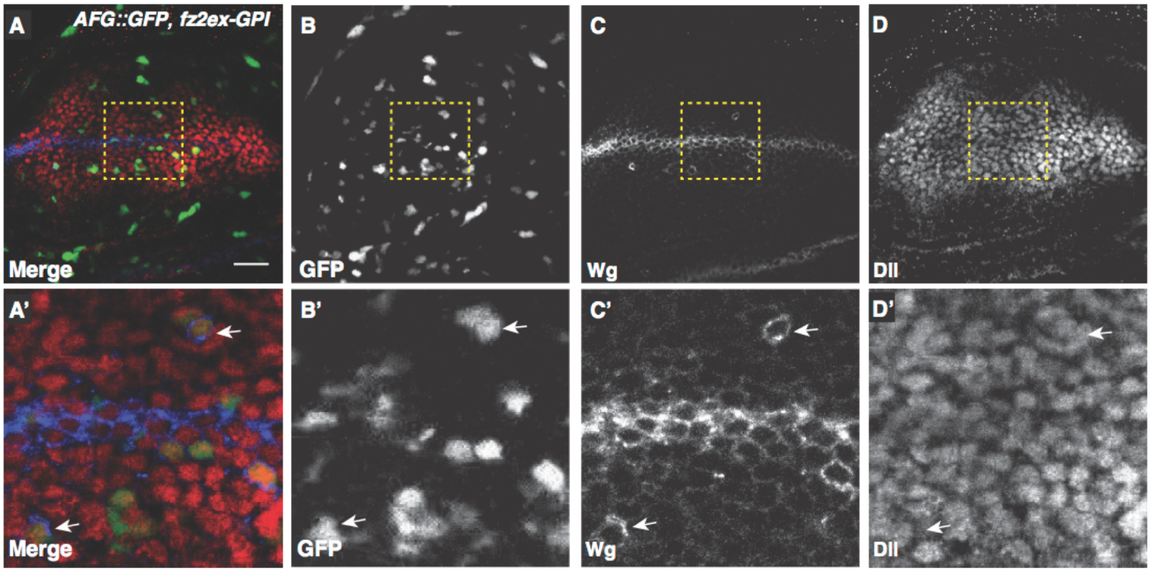


**Figure S2: Formation of a functional Wg-Fz2 complex is required for activation of Wg signalling at a distance.**

(A–D) Flip-out clones overexpressing dominant-negative Fz2 (marked with GFP). (A'–D') Area marked with yellow box in A–D. 24 hours after the heat-shock to induce DN-Fz2 expressing single-cell clones, total Wg staining shows clones with accumulation of Wg (C and C' arrows), however Dll levels were unaffected (D, D' arrows). Images show projection of 3 confocal slices. Scale bar 20 µm.

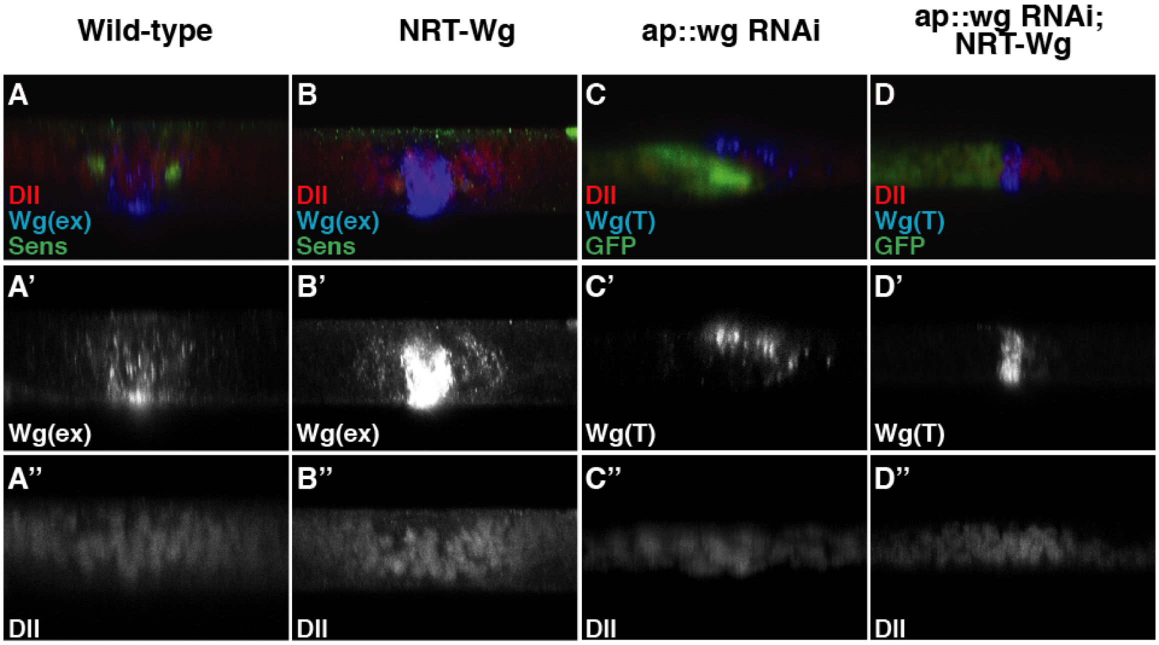


**Figure S3: Secreted Wg regulates target gene expression across the DV compartment boundary.**

(A-B) Cross-section view of extracellular Wg staining and Dll staining performed on either wild-type discs (A– A'') or homozygous *NRT-wg* discs and imaged with same confocal settings (B–B''). (C– D) Cross-section view of total Wg and Dll staining on discs with depletion of *wg* in dorsal compartment with *ap-Gal4* in either wild-type background (C–C'') or homozygous *NRT- wg* background (D–D''). a.u.= arbitrary unit.

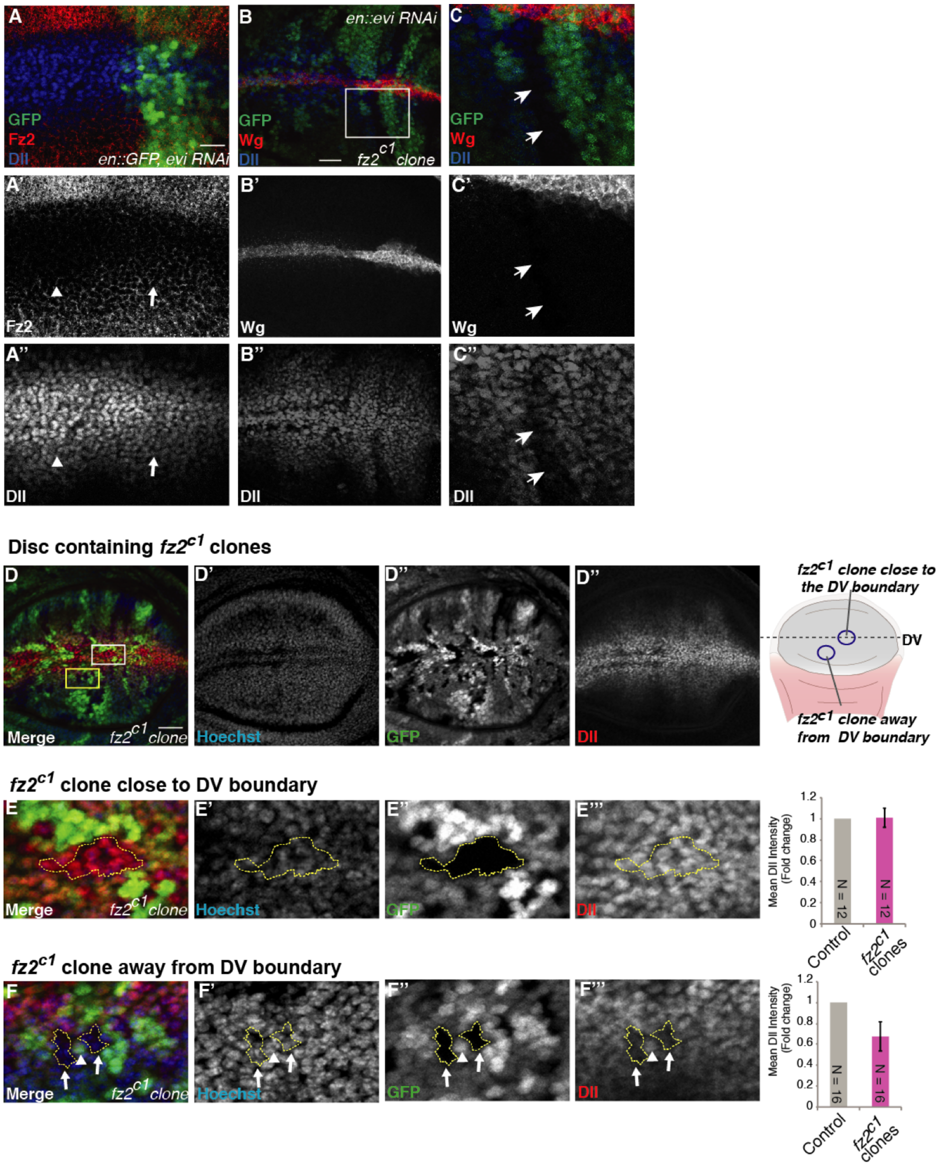


**Figure S4: Fz2 is required for the maintenance of long-range Wg target gene expression, related to Figure 4 -** (A–A'') Depletion of Evi in the posterior compartment of the disc shows up-regulation of Fz2 (compare arrow-head to arrow in A') while Dll levels are similar (compare arrowhead to arrow in A''). (B–B'') Depletion of Evi in the posterior compartment of the disc (identified with Wg accumulation) in the background of *fz2*$^{c1}$ clones (marked by absence of GFP) show reduced levels of Dll inside the mutant clones. (C–C'') Enlarged images of *fz2* mutant clone (marked by white box in B). (D–F) *fz2*$^{c1}$ clones in otherwise wild-type disc show no effect on Dll levels in clones close to the DV boundary (marked by white box in D). (E–E''') Enlarged images of the *fz2*$^{c1}$ clones found close to the DV boundary. (F'F'') Enlarged images of *fz2*$^{c1}$ found further away from the DV boundary (marked by yellow box in D) show reduced Dll levels inside the clones (F''' arrows) compared to control cells at the same distance (F''' arrowhead). Quantification shows Dll intensity measurement inside the clone and Dll intensity of the same area from neighbouring control cells. N represents combined number of clones from several discs. Scale bar 20 μm.

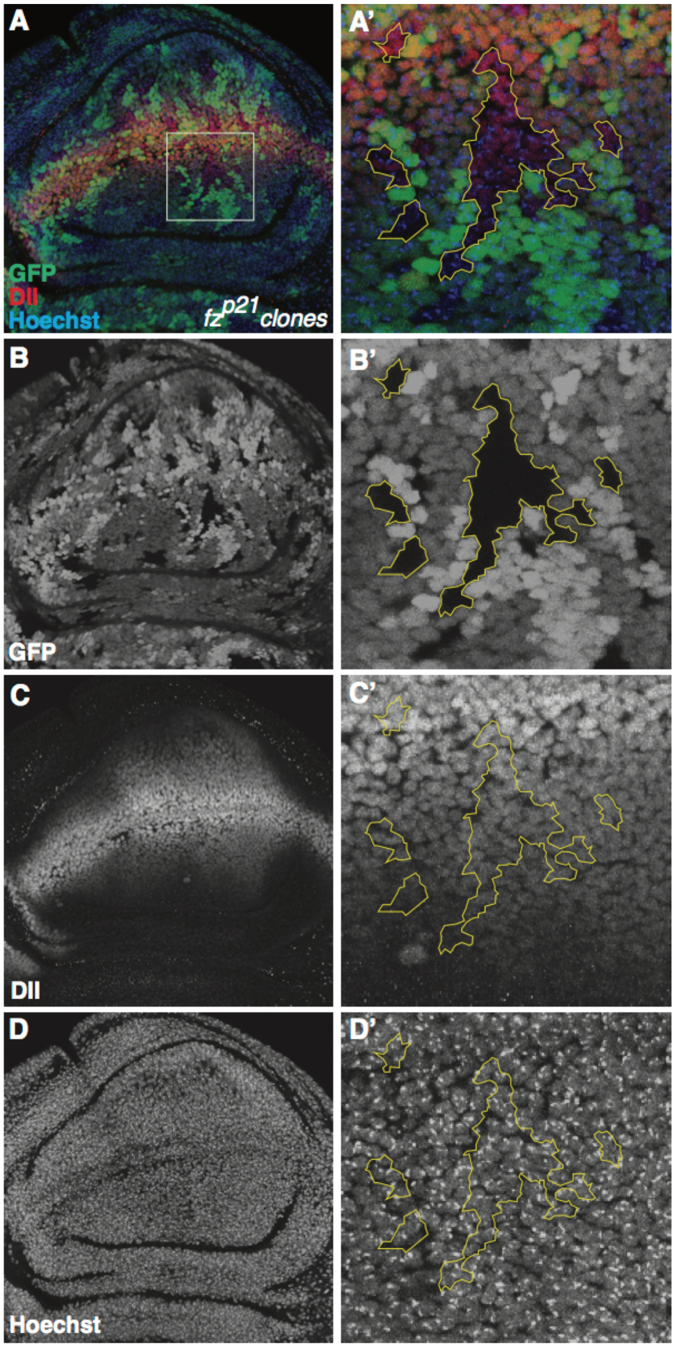


**Figure S5: Fz1 is not required for the maintenance of long-range Wg target gene expression, related to Figure 4 -** (A–D) *fz*$^{p21}$ clones in otherwise wild-type disc show no effect on Dll levels in clones. (A'–D') Enlarged images of the *fz*$^{p21}$ clones (marked by white box in A) show no significant difference in the Dll expression even at a distance from DV boundary. Scale bar 20 μm.
